# Stimulus familiarity and expectation jointly modulate neural activity in the visual ventral stream

**DOI:** 10.1101/192518

**Authors:** Mariya E. Manahova, Pim Mostert, Peter Kok, Jan-Mathijs Schoffelen, Floris P. de Lange

**Author notes:** **Corresponding author:** Mariya E. Manahova, Donders Institute for Brain, Cognition and Behaviour, Radboud University Nijmegen, P.O. Box 9101 6500 HB Nijmegen, Netherlands.

## Abstract

Prior knowledge about the visual world can change how a visual stimulus is processed. Two forms of prior knowledge are often distinguished: stimulus familiarity (i.e., whether a stimulus has been seen before) and stimulus expectation (i.e., whether a stimulus is expected to occur, based on the context). Neurophysiological studies in monkeys have shown suppression of spiking activity both for expected and for familiar items in object-selective inferotemporal cortex (IT). It is an open question, however, if and how these types of knowledge interact in their modulatory effects on the sensory response. In order to address this issue and to examine whether previous findings generalize to non-invasively measured neural activity in humans of both sexes, we separately manipulated stimulus familiarity and expectation, while non-invasively recording human brain activity using magnetoencephalography (MEG). We observed independent suppression of neural activity by familiarity and expectation, specifically in the lateral occipital complex (LOC), the putative human homologue of monkey IT. Familiarity also led to sharpened response dynamics, which was predominantly observed in early visual cortex. Together, these results show that distinct types of sensory knowledge jointly determine the amount of neural resources dedicated to object processing in the visual ventral stream.

## Significance Statement

The visual system is sensitive to whether it has encountered an item before or not, as well as whether an item is expected or surprising. The current study shows how these two factors jointly modulate the sensory response in the visual ventral stream: familiarity and expectation reduce the amplitude of the neural signal in object-selective visual cortex, while sharpening response dynamics in early visual cortex. Together, these mechanisms may aid efficient coding of visual input.

## Introduction

Our visual environment is complex and rapidly changing, making visual perception a challenging task. In order to quickly parse visual input and deliver a stable percept of the world, the visual system is thought to employ principles of operation that allow an efficient sensory representation of the most likely current state of the visual world (Rao and Ballard, 1999; Lee and Mumford, 2003; Friston, 2005; Meyer and Olson, 2011; Summerfield and de Lange, 2014). By consequence, the amount of experience the visual system has had with a particular object can influence how much resources are allotted to processing that object. For instance, viewing an object image repeatedly results in reduced spiking activity in inferior temporal (IT) cortex in monkeys (Miller et al., 1991) and reduced hemodynamic activity in the human homologue (Grill-Spector et al., 2006), lateral occipital cortex (LOC), as measured with functional magnetic resonance imaging (fMRI). These findings suggest that familiar items require fewer neural resources than unfamiliar items.

Structure in visual information can also affect visual processing. If images are regularly presented in a specific temporal sequence, it becomes possible to predict which image will be presented next. Studies find that expected object images elicit reduced spiking activity compared to unexpected items in monkey IT (Meyer and Olson, 2011; Kaposvari et al., 2016), while evidence is more mixed in human studies (Puri et al., 2009; Egner et al., 2010; Turk-Browne et al., 2010; Davis and Hasson, 2016), possibly due to differences in task demands (St. John-Saaltink et al., 2015).

Recently, Meyer et al. (2014) observed that image familiarization does not only lead to an activity reduction but also results in sharpening of the dynamics of neuronal visual responses in monkey IT: the sensory response to familiar images was truncated, leaving IT neurons in a state of readiness for ensuing images and thereby enhancing their ability to track rapidly changing displays. A similar temporal truncation has been seen for expected, compared to unexpected, images (Meyer and Olson, 2011). Temporal sharpening may complement neural activity suppression in representing the visual world in a maximally efficient manner.

While the effects of familiarity and expectation on the sensory response in monkey IT are relatively well described separately, it is uncertain whether and how these modulatory factors interact. Familiarity and expectation have to date been examined in distinct experimental paradigms, but since the two tend to go together (when we see a familiar image repeatedly, we come to expect it), their effects are easily confounded. Moreover, these processes have not been investigated extensively in humans using electrophysiological measures, and evidence from non-invasive recordings in humans is mixed. It is important to note that there is a nested relationship between familiarity and expectation, such that we must be familiar with an object before we can expect to see it. Because of this, expected and unexpected items are by necessity familiar, but, crucially, expected and unexpected items are equally familiar.

In the current study, we set out to study how image familiarity and expectation jointly determine the sensory response to object stimuli in humans by manipulating familiarity and expectation separately. We measured neural activity using magnetoencephalography (MEG) while participants viewed images that were familiar or novel and expected or unexpected. To preview, we found a reduction of activity for familiar compared to novel images in LOC. Within the class of familiar items, there was a further reduction of activity for expected compared to unexpected images in LOC. Moreover, we found a sharpening of response dynamics for familiar compared to novel images that was most prominent in early visual areas. These results show how familiarity and expectation independently modulate activity in object-selective visual cortex, possibly allowing efficient coding of visual input.

## Materials and Methods

### Participants

Twenty-nine healthy human volunteers (15 female, 14 male, mean age = 24.17 years, SD = 3.80 years) with normal or corrected-to-normal vision, recruited from the university’s participant pool, completed the experiment and received either monetary compensation or study credits. The sample size, which was defined a priori, ensured at least 80% power to detect within-subject experimental effects with an effect size of Cohen’s d>0.60. The study was approved by the local ethics committee (CMO Arnhem-Nijmegen, Radboud University Medical Center) under the general ethics approval (“Imaging Human Cognition”, CMO 2014/288), and the experiment was conducted in compliance with these guidelines. Written informed consent was obtained from each individual.

### Stimuli and Apparatus

#### Stimuli

Stimuli were chosen from the image set provided at http://cvcl.mit.edu/MM/uniqueObjects.html. A different object was represented in each image, and all objects were shown against a gray background. A total of 2054 images were presented for each participant. Familiar images were randomly selected for each pair of participants. Each pair of participants saw different familiar images, and the predictable and unpredictable images were counterbalanced within a pair of participants. Specifically, if for participant 1, set A comprised the predictable images and set B comprised the unpredictable images, the opposite was true for participant 2: set B comprised the predictable images, and set A comprised the unpredictable images. In both the behavioral and MEG sessions, the images subtended four degrees of visual angle.

#### Apparatus

MATLAB (The Mathworks, Inc., Natick, Massachusetts, United States) and the Psychophysics Toolbox extensions (Brainard, 1997) were used to show the stimuli on a monitor with a resolution of 1920x1080 pixels and a refresh rate of 100 Hz. For the MEG session, a PROpixx projector (VPixx Technologies, Saint-Bruno, QC Canada) was used to project the images on the screen, with a resolution of 1920x1080 and a refresh rate of 100 Hz.

### Trial structure

For the behavioral training session as well as for the MEG testing session, each trial began with a fixation dot (see Figure 1A for the trial structure). The fixation dot was presented for a randomly selected period between 500 and 750 ms. Then, six images were shown, each lasting for 180 ms and presented back-to-back. At the end of a trial, if an oddball was presented during the trial and a response was given, the fixation dot turned green for 500 ms. If the response was incorrect, the fixation dot turned red for 500 ms. A response was considered incorrect on three occasions: if the participant pressed the button during a trial with an oddball stimulus but before the oddball was presented; if the participant pressed the button on a trial where no oddball was presented; or if the participant did not press the button on a trial where an oddball was presented. If no oddball was presented and no response was given, the fixation dot did not change color and the white-and-black fixation dot remained on the screen for 750 ms. At the end of each trial, a blank screen was presented for 1250 ms, and participants were encouraged to blink during this period.

**Figure 1.**
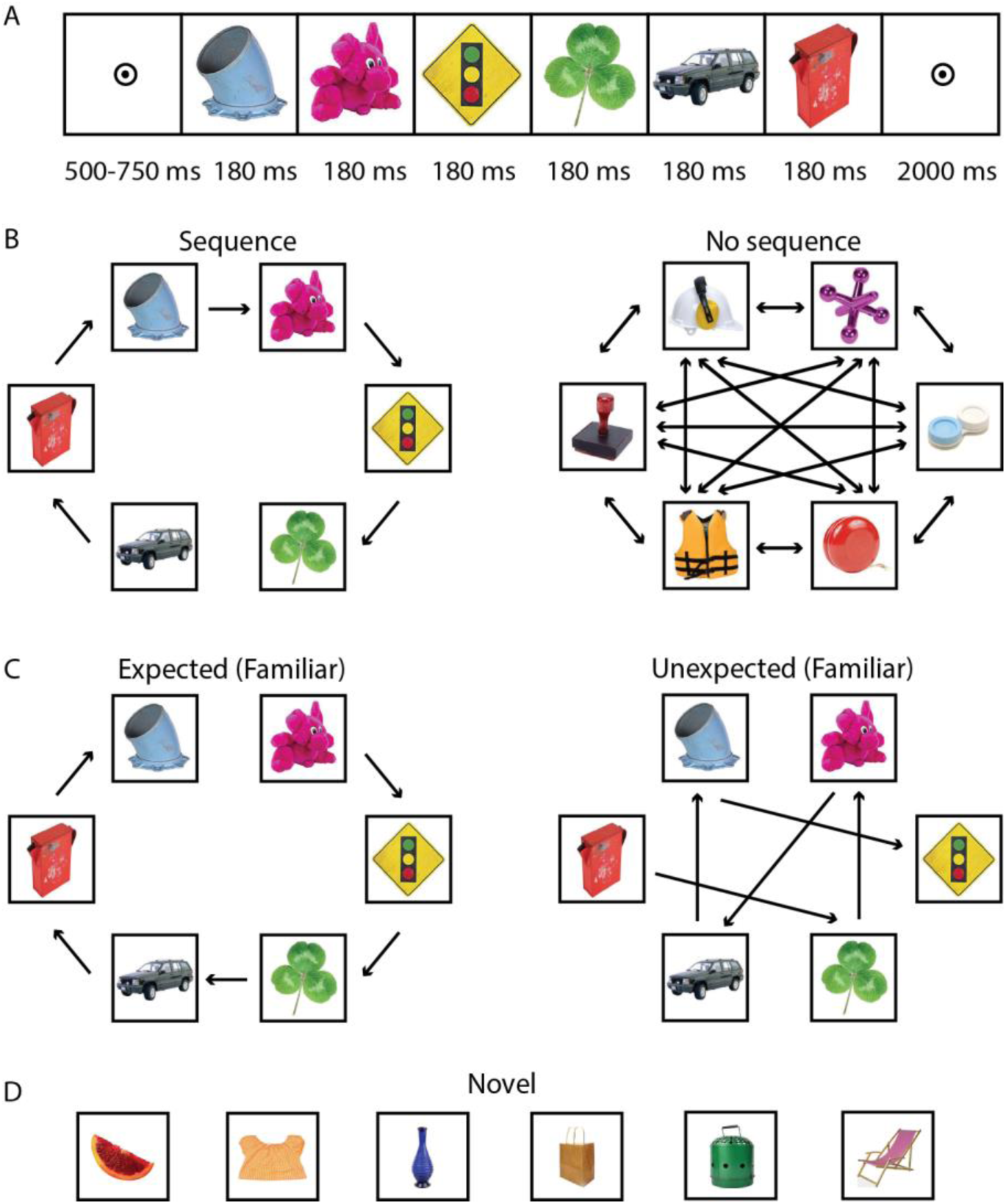
(A) Trial structure. First, participants saw a bull’s eye fixation point for a jittered period between 500 and 750 ms. Then, each of the six images was presented for 180 ms with no gap between images. Finally, a fixation dot was presented at the end of the trial. (B, C, D) Experimental design: (B) In one condition, images were always presented in a specific circular sequence (left). In another condition, images were presented in any sequence (right). (C) For the condition with a specific sequence, images were sometimes presented in the expected order (left), while on other occasions images were presented in an unexpected order (right). (D) New images were presented during novel trials, and each unique image was only shown once during the whole experiment.

### Experimental Design and Statistical Analysis

#### Experimental Design

First, participants completed a behavioral training session in which they were familiarized with two sets of six images, one set with predictable structure and one set with no predictable structure. More specifically, they observed the predictable images always in the same order, while the unpredictable images were shown in a randomly shuffled order on each trial (Figure 1B). Importantly, the order for the predictable images was circular, i.e., each of the six images could be presented first. Predictable images comprised 50% of trials, and unpredictable images comprised the other 50%. Participants performed an oddball detection task by pressing the spacebar when they saw an image of a rubber duck. Images of duckies were presented on 10% of trials as one of the six images in the sequence. The duckies were of eight different colors and there were two viewpoints per color for a total of sixteen images of duckies. Multiple images of duckies were used to reduce the possibility that participants may attend selectively to a particular color (e.g., yellow) or shape. The oddball task was chosen such that participants were required to maintain their attention on the visual stream.

During the behavioral training session, participants completed 10 blocks of 80 trials each for a total of 800 trials. Each block lasted 4.9 minutes, leading to a total training session duration of approximately one hour. At the end of the behavioral training session, participants’ knowledge of the order of predictable images was assessed with a sequence identification task. Participants were shown one of the six images from the sequenced set, and they had to indicate which of the other five was most likely to follow it. This was done for each of the six images in the predictable set. The assessment took about three minutes.

One or two days later, participants completed the MEG testing session in which they saw familiar (i.e., those presented during the behavioral session) and novel (never seen before) images (see Figure 1B, C, and D for a depiction of the conditions). In contrast to the training session, the predictable images were now sometimes presented in the learned order (expected, 50% of sequenced trials) and sometimes in a shuffled order (unexpected, 50% of sequenced trials). The shuffled sequences for unexpected trials were chosen in such a way that each image in the sequence was followed by an unpredicted image; in other words, none of the images were followed by the image they predicted (see Figure 1C). The unpredictable images were shown in shuffled orders, as during the training session. Unpredictable images comprised one third of trials, and predictable images also comprised one third of trials. The remaining third of trials was comprised of novel images that the participants had not seen before (see Figure 1D). Unique novel images were used for every trial, so each novel image was only shown once during the experiment. Participants performed the same oddball task as during the training session: they had to respond when they saw a ducky, and duckies were presented on 10% of trials. During the MEG testing session, participants completed 8 blocks of 120 trials each for a total of 960 trials. Each block lasted 7.4 minutes, leading to a total experimental duration of approximately one hour. At the end of the MEG testing session, participants’ knowledge of the familiar images was assessed. Participants saw 60 images, the twelve familiar ones and 48 selected at random from the novel images participants had been shown, and participants had to indicate whether the image was familiar or novel. ‘Familiar’ referred to images seen repeatedly during the behavioral training session as well as during the MEG testing session, while ‘novel’ referred to images seen only once during the MEG testing session.

#### Statistical analysis

For the behavioral results, mean reaction time and accuracy were first calculated within participant per condition. Then, two-tailed paired-samples t-tests were calculated for the two relevant conditions for a comparison. Behavioral data were analyzed for 28 out of the 29 participants, as a technical issue in data acquisition prevented the analysis of behavioral data of the first participant.

In order to statistically assess the MEG activity difference between conditions in the time domain and control for multiple comparisons, we applied cluster-based permutation tests (Maris and Oostenveld, 2007), as implemented by FieldTrip (Oostenveld et al., 2011). The tests were carried out on the time period between 0 and 1200 ms, 0 ms being the onset of the first stimulus, over all sensors, and 10,000 permutations were used per contrast. For each sensor and time point, the MEG signal was compared univariately between two conditions, using a paired *t*-test. Positive and negative clusters were then formed separately by grouping spatially and temporally adjacent data points whose corresponding *p*-values were lower than 0.05 (two-tailed). Cluster-level statistics were calculated by summing the *t*-values within a cluster, and a permutation distribution of this cluster-level test statistic was computed. The null hypothesis was rejected if the largest cluster in the considered data was found to be significant, which was the case if the cluster’s *p*-value was smaller than 0.05 as referenced to the permutation distribution. The standard error of the mean was computed using a correction that makes it suitable for within-subject comparisons (Cousineau, 2005; Morey, 2008).

We also applied cluster-based permutation tests in order to statistically assess MEG activity differences between conditions in the frequency domain (see *Data Analysis*). The tests were carried out on the log_10_-transformed data for the frequency of interest (stimulus presentation frequency: 5.6 Hz), over all sensors, and with 10,000 permutations per contrast. Adjacent sensors with nominal *p*-values lower than 0.05 (two-tailed) were grouped into clusters. The *t*-values within a cluster were summed, yielding a cluster-level statistic. If the largest cluster’s *p*-value was smaller than 0.05, the difference across the compared conditions was considered statistically significant.

To determine whether our data offered evidence for surprise enhancement or expectation suppression (or both), we compared familiar unexpected trials with unpredictable trials, as well as familiar expected with unpredictable trials. For these comparisons, the time domain data were averaged per condition over a time period from 500 until 900 ms after the onset of the visual sequence, since this was previously identified as the time period of the largest difference between expected and unexpected stimulus streams. Next, two paired-samples t-tests were carried out, to statistically compare the amplitude of unexpected and unpredictable trials, as well as the amplitude of expected and unpredictable trials.

Finally, in order to assess whether explicit knowledge of the sequence immediately after training influenced the activity modulation by expectation, we divided participants based on whether their performance was above or below chance level (20%) on the sequence identification task. This division resulted in two groups of 14 participants each. We averaged each participant’s data per condition from 500 ms to 900 ms and over the sensors that contributed to the significant difference for expected vs. unexpected sequences. We examined whether there were differences between the groups using an independent-samples t-test.

### Data acquisition

#### MEG recordings

Brain activity was recorded using a 275-channel MEG system with axial gradiometers (VSM/CTF Systems, Coquitlam, BC, Canada) in a magnetically shielded room. During the experiment, head position was monitored online and corrected if necessary (Stolk et al., 2013). This method uses three coils: one placed on the nasion, one in an earplug in the left ear, and one in an earplug in the right ear. To aid in the removal of eye- and heart-related artifacts, horizontal and vertical electrooculograms (EOGs) as well as an electrocardiogram (ECG) were recorded. A reference electrode was placed on the left mastoid. The sampling rate for all signals was 1200 Hz. A projector outside the magnetically shielded room projected the visual stimuli onto a screen in front of the participant via mirrors. Participants gave their behavioral responses via an MEG-compatible button box. Participants’ eye movements and blinks were also monitored using an eye-tracker system (EyeLink, SR Research Ltd., Mississauga, Ontario, Canada).

#### MRI Recordings

To allow for source reconstruction, anatomical magnetic resonance imaging (MRI) scans were acquired using a 3T MRI system (Siemens, Erlangen, Germany) and a T1-weighted MP-RAGE sequence with a GRAPPA acceleration factor of 2 (TR = 2300 ms, TE = 3.03 ms, voxel size 1 × 1 × 1 mm, 192 transversal slices, 8° flip angle).

### Data analysis

#### MEG data analysis

The MEG data were preprocessed offline using the FieldTrip software (Oostenveld et al., 2011). Trials where oddball stimuli were presented and/or a response was given were removed from analysis. This was done because oddballs and responses elicited neural activity unrelated to the research question. Then, trials with high variance were manually inspected and removed if they contained excessive and irregular artifacts. Afterwards, independent component analysis (ICA) was applied to identify regular artifacts such as heartbeat and eye blinks. The independent components for each participant were then correlated to the horizontal and vertical EOG signals and to the ECG signal. In this way, it was possible to identify which components most likely corresponded to the heartbeat and eye blinks.

#### Event-related fields

Before calculating event-related fields (ERFs), a low-pass filter at 30 Hz was applied to the data. The data were baseline-corrected on the interval starting at 200 ms before stimulus onset until stimulus onset (0 ms). Subsequently, the data were transformed to simulate planar gradiometers in order to facilitate interpretation as well as averaging over participants. We took an equal number of trials per condition for each comparison to avoid any possible confounding influence of noise due to unequal number of trials. We did this by choosing a random selection of trials from the condition with more trials to match the number of trials in the condition with fewer trials.

#### Source reconstruction on time-domain data

We performed source reconstruction in order to facilitate interpretation of the ERFs. Source reconstruction was done for 27 of the participants for whom we were able to acquire a structural MRI scan. We created volume conduction models, based on a single shell model of the inner surface of the skull, and subject-specific dipole grids, which were based on a regularly spaced 6-mm grid in normalized MNI (Montreal Neurological Institute) space. For each grid point the lead fields were rank reduced by removing the sensitivity to the direction perpendicular to the surface of the volume conduction model. Source activity was then obtained by estimating linearly constrained minimum variance (LCMV) spatial filters (van Veen et al., 1997), for which the data covariance was calculated over the interval of 200-1200 ms post-stimulus and regularized using shrinkage (as described in Blankertz et al., 2011) with a regularization parameter of 0.01. The filters were applied to the axial gradiometer data and resulted in an estimated two-dimensional dipole moment per grid point, per time point.

For visualization as well as interpretation, we reduced these two-dimensional moments to a scalar value by taking the norm of the vector. This value reflects the degree to which a particular source location contributes to (differences in) activity measured at the sensor level. Critically, this value was obtained from the difference ERF between two conditions, rather than from each condition individually and subtracted afterwards. In this way, differences in dipole orientation are also captured, instead of only magnitude, which would presumably correspond to different neural populations within the same source location.

One problem with taking the norm of the vector is that this is always a positive value and will therefore, due to noise, suffer from a positive bias. To counter this bias, we employed a permutation procedure, in which the condition labels were shuffled across trials. A total of 1000 permutations were performed, the average of which was taken as an estimate of the noise. Specifically, the average was calculated over the square of the dipole’s norm (i.e., after squaring and summing in the Pythagorean theorem but before taking the square root). Next, this noise estimate was subtracted from the (square of) the true data, after which the data were divided by the noise estimate in order to counter the depth bias. The resulting values were then averaged over participants and negative values were set to zero. Finally, the square root was taken, resulting in a group-level estimate of the contributions of each source location.

#### Spectral analysis

We conducted a spectral analysis for all frequencies between 1 and 30 Hz. We applied the fast Fourier transform to the planar-transformed time domain data, after tapering with a Hanning window. The time period of interest was from 180 until 1080 ms, and data were baseline-corrected on the interval starting at 200 ms before stimulus onset until stimulus onset (0 ms). The spectral analysis was carried out separately per condition, and the resulting power per frequency was averaged over participants.

#### Source reconstruction on frequency-domain data

We also applied source reconstruction analysis in order to facilitate interpretation of the power spectra. The source models and lead fields were obtained as described before and for the same 27 participants. Source activity was obtained by applying spatial filters based on partial canonical correlations (PCC; Schoffelen et al., 2008) from the power data described above. The PCC method allows for the efficient extraction of the source-level power for single trials. The regularization parameter was 0.01, and the frequency of interest was 5.6 Hz. This procedure resulted in an estimated three-dimensional dipole moment per grid point. For each grid point, we calculated the mean across each of the three spatial dimensions, computed its absolute value, and squared it. Then, we summed the resulting values for the three dimensions, which produced a single value per grid point. This analysis was carried out separately for each condition, and afterwards we averaged the resulting values across participants.

## Results

### Behavioral results

The participants’ task was to press a button whenever they saw an oddball stimulus, in this case an image of a ducky. Participants were at near ceiling level in their performance on the oddball task (mean accuracy = 94.9%, SD = 2.9%). Participants’ accuracy was not significantly affected by whether duckies appeared in familiar vs. novel sequences (t_27_ = −0.75, *p* = 0.46), or in expected vs. unexpected sequences (t_27_ = 0.40, *p* = 0.69). Furthermore, participants’ reaction times to oddball trials were not significantly different between duckies embedded in familiar vs. novel sequences (t_27_= −1.74, *p* = 0.09) or expected vs. unexpected sequences (t_27_ = −0.04, *p* = 0.97).

At the end of the behavioral training session, participants’ knowledge of the order of the predictable images was assessed. On average, when participants were shown an image and had to report which image followed it, they selected the correct image 25% of the time (SD = 19.7%), with chance level performance at 20%. This suggests that subjects were largely unaware of the sequence, in agreement with their verbal reports.

At the end of the MEG session, participants’ knowledge of image familiarity was assessed. On average, when participants had to report whether an image was familiar or novel, they correctly identified the familiar images in 91.9% of trials (SD = 5.8%), showing that they were clearly aware of the image familiarity manipulation.

### MEG results

#### Familiar items lead to reduced activity in LOC

To investigate the difference in amplitude between familiar and novel items without any influence of the expectation manipulation, we compared the familiar unpredictable and novel conditions since participants did not learn a sequence for the images in the unpredictable condition. A significant difference (*p* < 1e-6) across conditions was observed, which was driven by the cluster of sensors shown in Figure 2A from approximately 200 ms until 1200 ms. The black asterisks in the figure denote sensors that contribute to this cluster for at least half of the time period from 200 ms to 1200 ms. The average timecourse for the sensors contributing to the cluster is plotted in Figure 2C; the black line at the bottom shows that at least one of the selected sensors at that time point contributes to the significant difference. Clearly, novel items lead to significantly more activity than familiar ones. Source reconstruction revealed that the difference between familiar and novel items stemmed primarily from sources along the visual stream. This included early visual areas as well as downstream visual areas such as right and left inferior occipital gyrus (Figure 2B, Table 1), in the vicinity of lateral occipital complex (LOC) (Grill-Spector et al., 2000).

**Table 1.**
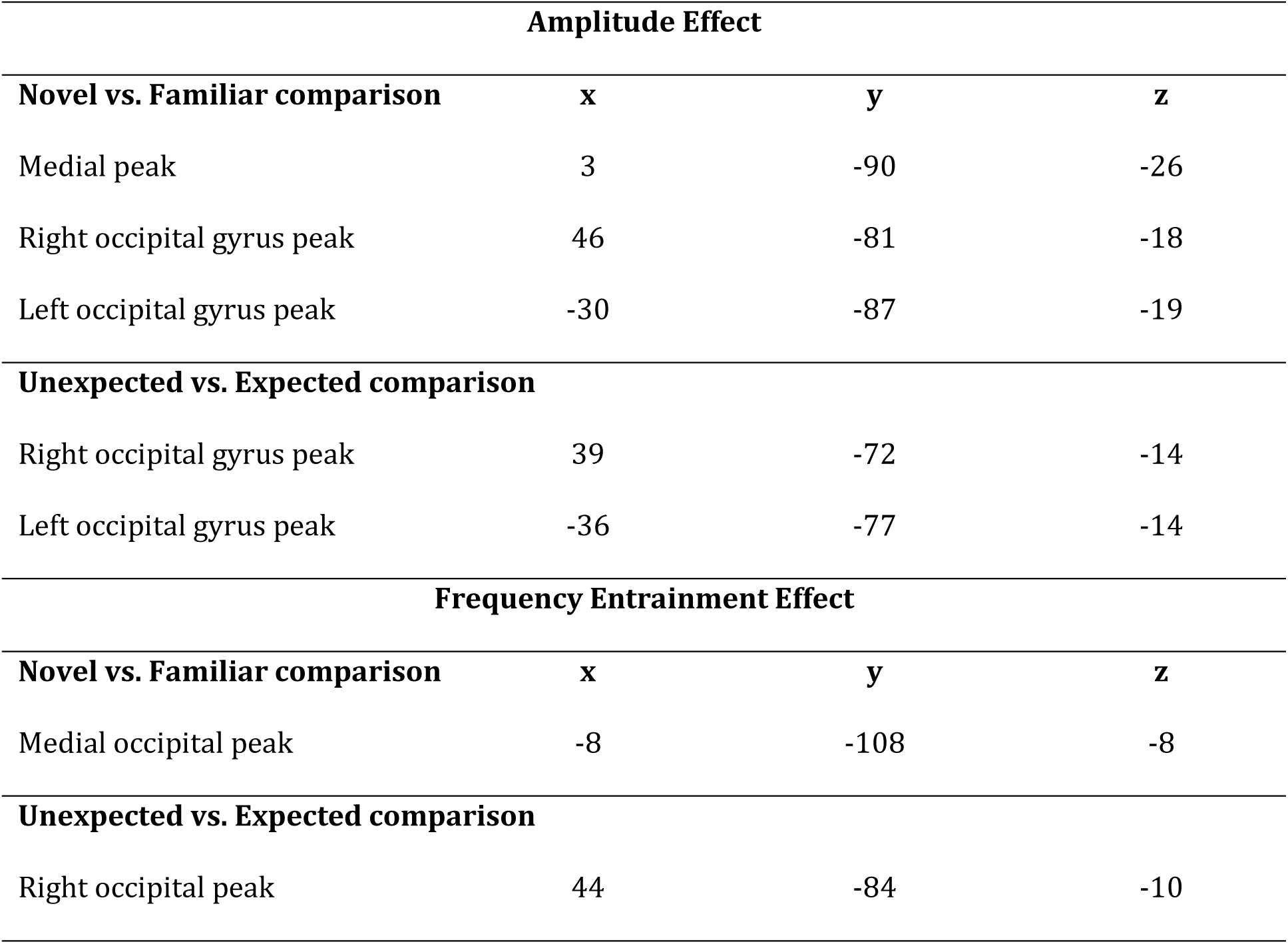
MNI coordinates of peak source locations for the respective comparisons. Reported values are mean values across participants in mm.

| **Amplitude Effect** |  |  |  |
| --- | --- | --- | --- |
| **Novel vs. Familiar comparison** | **x** | **y** | **z** |
| Medial peak | 3 | -90 | -26 |
| Right occipital gyrus peak | 46 | -81 | -18 |
| Left occipital gyrus peak | -30 | -87 | -19 |
| **Unexpected vs. Expected comparison** |  |  |  |
| Right occipital gyrus peak | 39 | -72 | -14 |
| Left occipital gyrus peak | -36 | -77 | -14 |
| **Frequency Entrainment Effect** |  |  |  |
| **Novel vs. Familiar comparison** | **x** | **y** | **z** |
| Medial occipital peak | -8 | -108 | -8 |
| **Unexpected vs. Expected comparison** |  |  |  |
| Right occipital peak | 44 | -84 | -10 |

**Figure 2.**
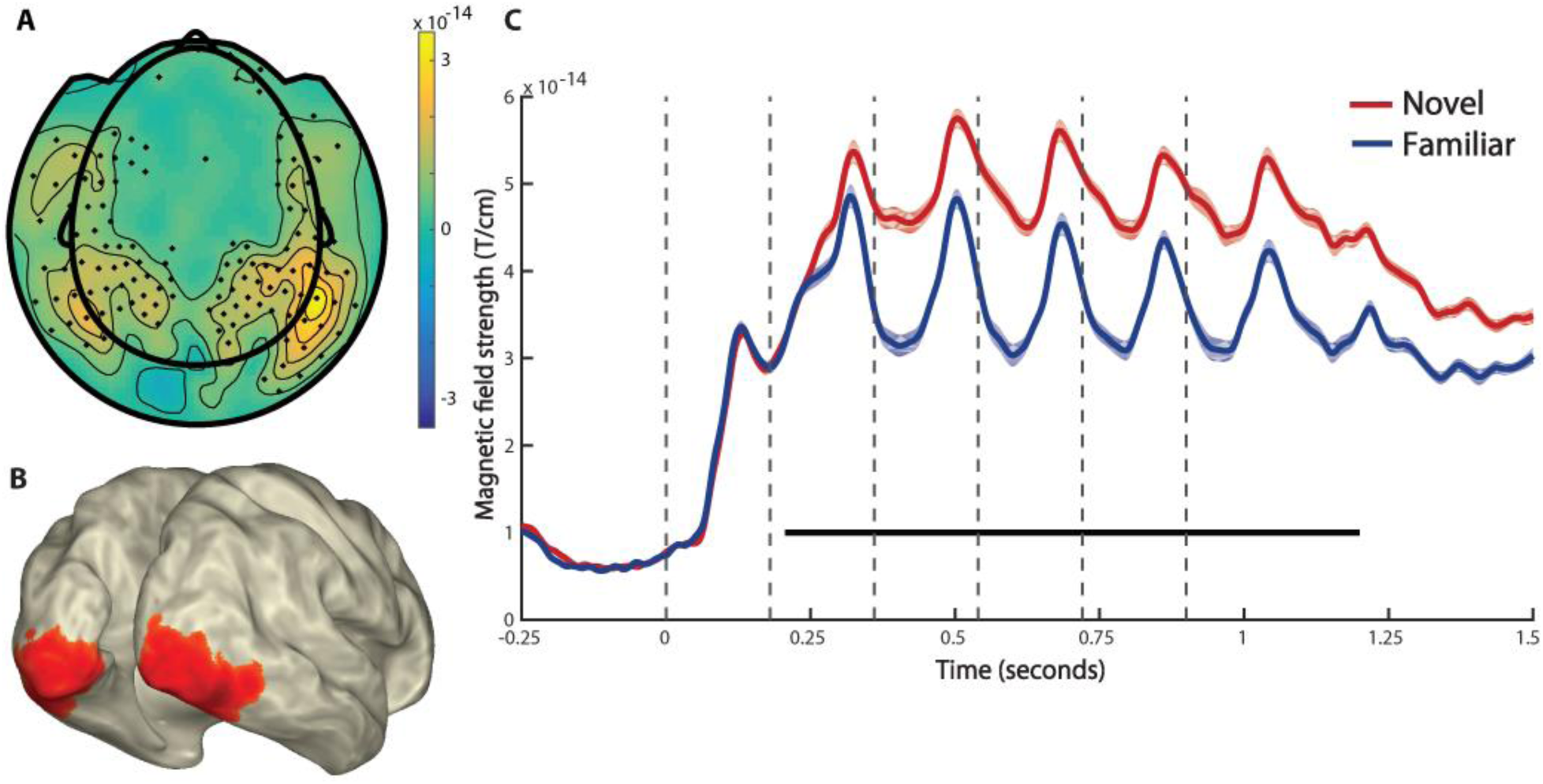
Familiarity effect on amplitude. (A) Topography of the difference in amplitude between the familiar unpredictable and novel conditions (novel – familiar). Black asterisks mark sensors that contribute to the significant cluster for at least half of the time period from 200 ms to 1200 ms. (B) Source reconstruction of familiar vs. novel. Activity was averaged over the time period of 200 to 1200 ms and interpolated onto a cortical surface. Plotted activity was thresholded at 80% of peak value for illustration purposes. (C) Activity over time for the familiar unpredictable (blue) and novel (red) conditions. Activity was averaged over sensors highlighted in (A). Shaded areas are error bars illustrating within-subject SEM for the familiar (light blue) and novel (light red) conditions. Horizontal black bar at the bottom shows that at least one of the selected sensors contributes to the significant cluster at this time point. Dotted vertical lines denote the onset of each image.

#### Unexpected items lead to enhanced activity in LOC

To examine the difference in amplitude between expected and unexpected items when familiarity was held constant, we compared the expected and unexpected conditions, both of which consisted of familiar images. A significant difference (*p* = 0.008) across conditions was observed, which was driven by the cluster of sensors shown in Figure 3A from approximately 500 ms until 900 ms. The black asterisks in the figure denote sensors that contribute to this cluster for at least half of the time period from 500 ms to 900 ms. The timecourse for the sensors contributing to the cluster is plotted in Figure 3C; the black line at the bottom shows that at least one of the selected sensors contributes to a significant difference at that time point. Evidently, unexpected items lead to significantly more activity than expected ones. Source analysis demonstrated that the difference between expected and unexpected items could be localized to the right inferior occipital gyrus and to a lesser degree to the left inferior occipital gyrus (Figure 3B, Table 1), corresponding to right and left LOC.

**Figure 3.**
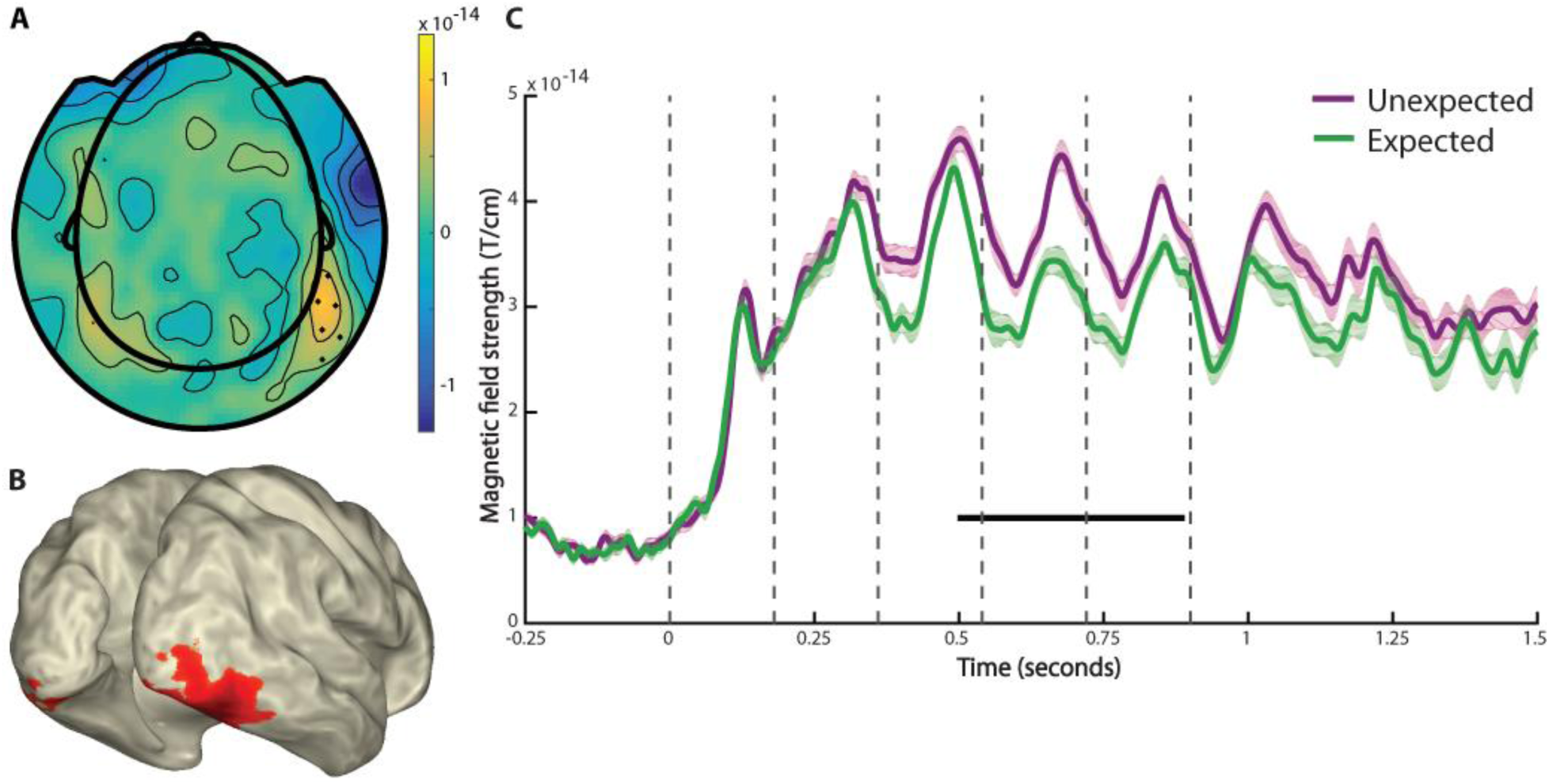
Expectation effect on amplitude. (A) Topography of the difference in amplitude between the expected and unexpected conditions (unexpected - expected). Black asterisks mark sensors that contribute to the significant cluster for at least half of the time period from 500 ms to 900 ms. (B) Source reconstruction of expected vs. unexpected. Activity was averaged over the time period of 500 to 900 ms and interpolated onto a cortical surface. Plotted activity was thresholded at 80% of peak value for illustration purposes. (C) Activity over time for the expected (green) and unexpected (purple) conditions. Activity was averaged over sensors highlighted in (A). Shaded areas are error bars illustrating within-subject SEM for the expected (light green) and unexpected (light purple) conditions. Horizontal black bar at the bottom shows that at least one of the selected sensors contributes to the significant cluster at this time point. Dotted vertical lines denote the onset of each image.

Moreover, we asked whether our data provided support for surprise enhancement or expectation suppression. To this end, we compared familiar unpredictable to both familiar unexpected and familiar expected items, respectively, since the former contrast illustrates the effect of a violated expectation and the latter demonstrates the effect of a confirmed expectation. Unexpected trials showed stronger activity than familiar unpredictable trials (t_28_ = −3.01, *p* = 0.006), demonstrating an activity increase for violated expectations. Expected trials, on the other hand, did not show a robust difference to familiar unpredictable trials (t_28_ = −1.53, *p* = 0.14).

#### Familiarity sharpens the dynamics of the sensory response

Sharper response dynamics include a truncated sensory response, which leads to higher dynamic range (peak-to-trough excursion) of the response (Meyer et al., 2014). The dynamic range of the signal can be approximated by the power at the driving frequency. To investigate the difference in power at the stimulus frequency, we compared the novel and familiar conditions. A significant difference (*p* = 0.016) emerged for the driving frequency of 5.6 Hz in the cluster of sensors shown in Figure 4A. The black asterisks in the figure denote sensors that contribute to the significant cluster, and the power spectrum for these sensors is plotted in Figure 4C. Familiar items led to significantly more power at the stimulus frequency of 5.6 Hz than novel items. Source reconstruction revealed that the largest difference in power between the familiar and novel conditions was in a medial posterior location belonging to early occipital cortex (Figure 4B, Table 1).

**Figure 4.**
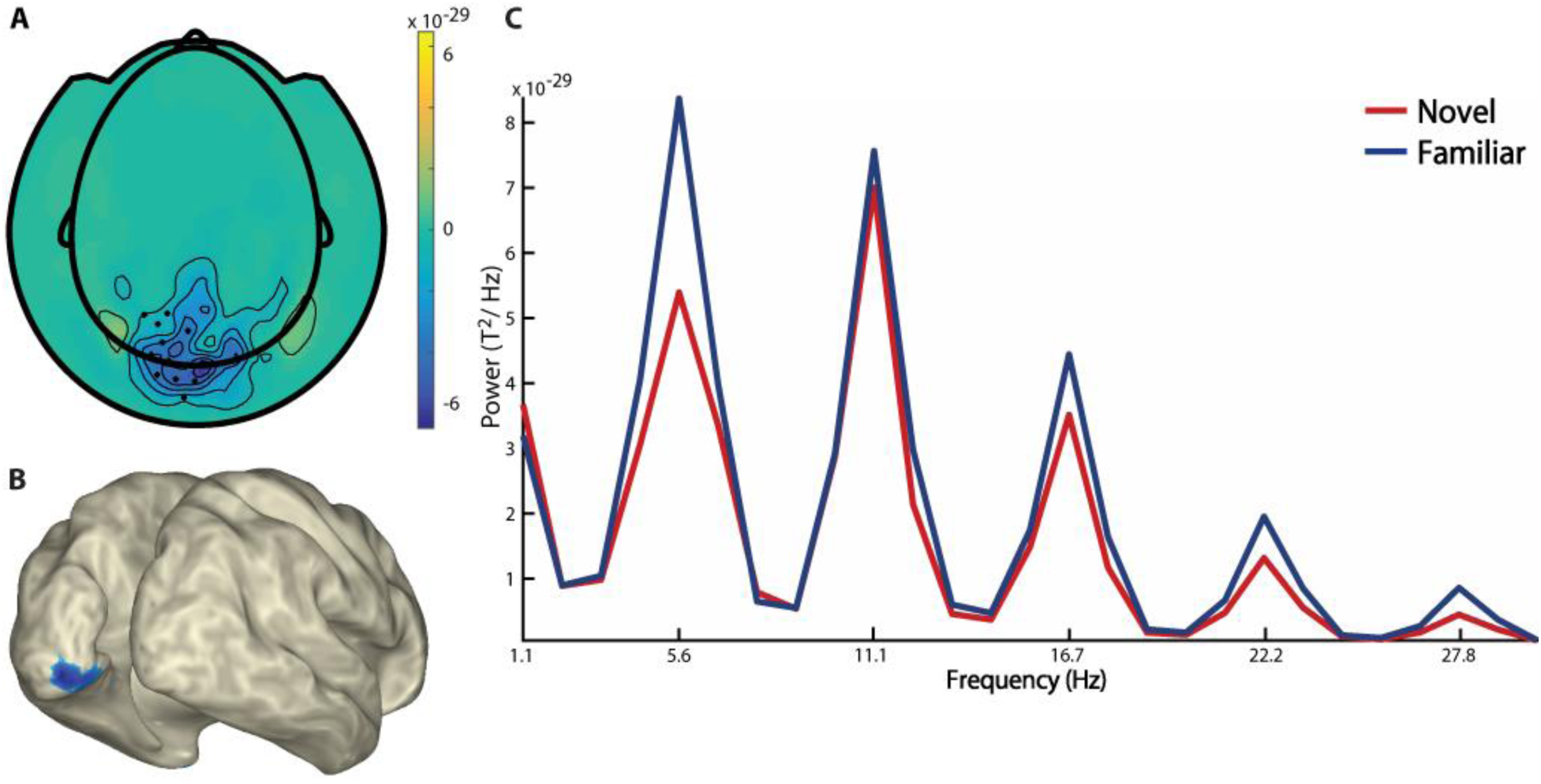
Familiarity effect on power. (A) Topography of the difference in power between the familiar unpredictable and novel conditions (novel – familiar). Black asterisks mark sensors that contribute to the significant cluster for the 5.6 Hz frequency. (B) Source reconstruction of familiar vs. novel. Activity corresponding to the frequency of 5.6 Hz was interpolated onto a cortical surface. Plotted activity was thresholded at 50% of peak value for illustration purposes. (C) Power spectrum of familiar unpredictable (blue) and novel (red). Activity was averaged over sensors highlighted in (A).

#### Expectation does not significantly sharpen the dynamics of the sensory response

To examine whether expectation likewise sharpened the neural response dynamics, we compared the difference in power at the stimulus frequency between the unexpected and expected conditions. There was no significant difference (*p* = 0.700) for the driving frequency of 5.6 Hz, as shown in Figure 5A. The power spectrum for all sensors is plotted in Figure 5C. It suggests that expected items may lead to more power at the stimulus frequency of 5.6 Hz than unexpected items, but this difference was not significant. Source analysis demonstrated that the largest difference in power between the expected and unexpected conditions was in early visual areas (Figure 5B, Table 1), but it is difficult to interpret this outcome since the effect is not statistically significant.

**Figure 5.**
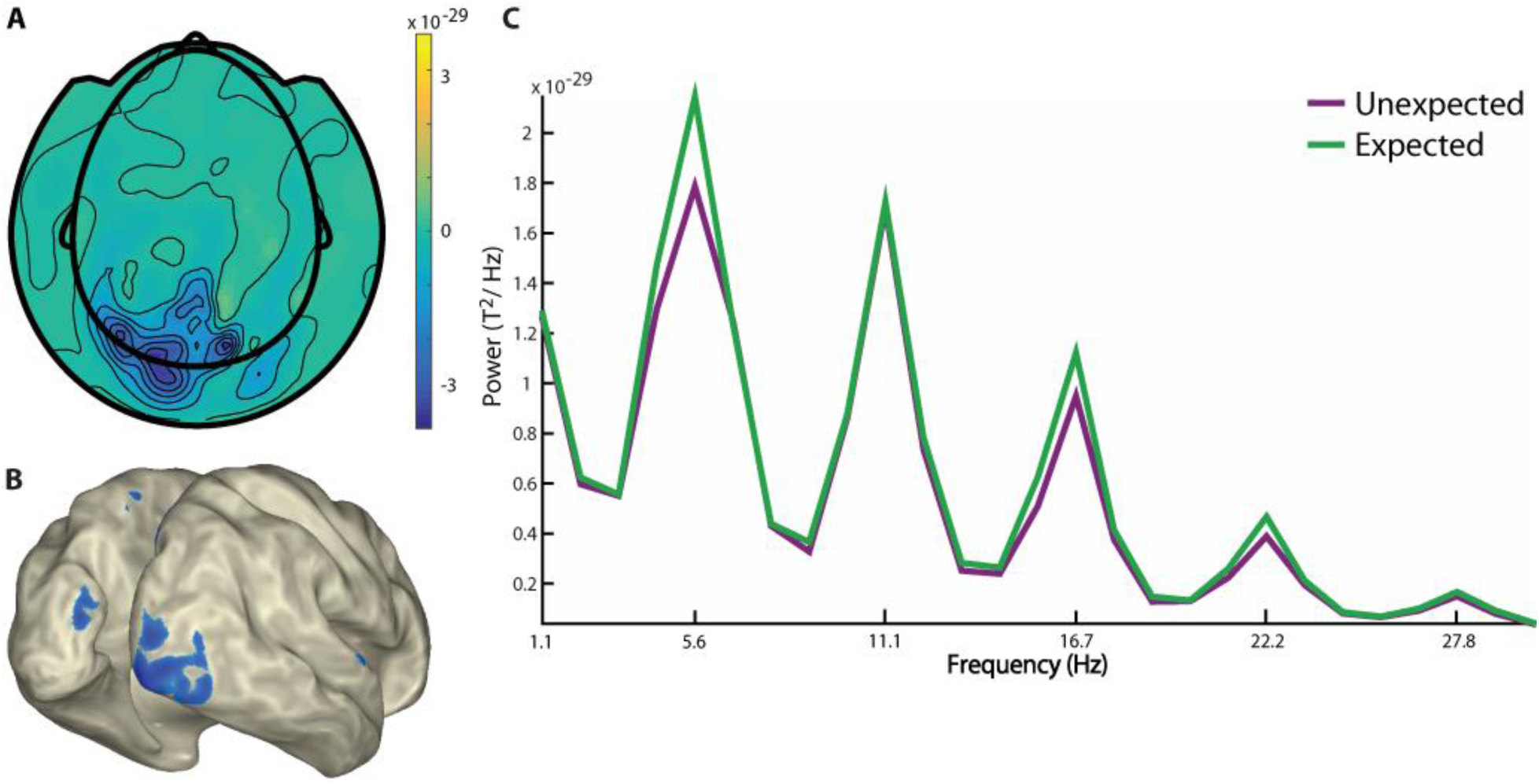
Expectation effect on power. (A) Topography of the difference in power between the expected and unexpected conditions (unexpected - expected). The lack of black asterisks denotes that no sensors contribute to a cluster for the 5.6 Hz frequency. (B) Source reconstruction of expected vs. unexpected. Activity corresponding to the frequency of 5.6 Hz was interpolated onto a cortical surface. Plotted activity was thresholded at 50% of peak value for illustration purposes. (C) Power spectrum of expected (green) and unexpected (purple). Activity was averaged over all sensors.

#### Explicit knowledge of sequence immediately after training does not modulate the expectation effect

We wanted to determine whether the extent to which participants could report the order of the stimuli influenced the expectation effect. Thus, we compared the expectation effect in the group which performed above chance on the sequence identification task to that in the group performing below chance. Although expectation modulated the amplitude of MEG activity in each group separately (t_13_ = −3.28, *p* = 0.006 for the above-chance group and t_13_ = −2.83, *p* = 0.01 for the below-chance group), this modulation did not differ significantly between the two groups (t_26_ = −0.27, *p* = 0.79). This suggests that the effect of expectation on the magnitude of the neural response is not related to participants’ explicit knowledge of the stimulus structure.

## Discussion

The sensory response to a stimulus can be modulated by whether a stimulus has been seen before (i.e., stimulus familiarity), as well as by whether a stimulus is expected or unexpected in the current context (i.e., stimulus expectation). In this study, we manipulated stimulus familiarity and expectation separately, which allowed us to examine the independent and shared effects of familiarity and expectation on brain activity in the visual system, using MEG. We found that familiar images elicited markedly less neural activity than novel images in early visual and object-selective lateral occipital (LOC) cortex. Similarly, expected images were also associated with reduced neural activity compared to unexpected images in LOC. The independent manipulation of familiarity and expectation in our study allows us to conclude that these distinct types of sensory knowledge jointly determine the amount of neural resources dedicated to object processing in the visual ventral stream.

Familiar items, when compared to novel items, were also associated with a temporally truncated (i.e., temporally sharpened) sensory response. In the context of our paradigm, in which stimuli rapidly followed each other, this led to an increased dynamic range of the signal. This was visible as increased power in the stimulus frequency, which was most prominent in early visual cortex. A similar trend, albeit non-significant, was present for expected vs. unexpected items. The sharper response dynamics for familiar than novel stimuli we observed replicate earlier findings by Meyer et al. (2014), although we observe the strongest contribution in early, rather than later, visual regions. While an increase in attention can also lead to an increase of power in the stimulus frequency for visually entrained stimuli (Ding et al., 2006), it is unlikely that this underlies the sharpened response for familiar items, for two reasons. Firstly, given that participants’ task was to detect oddball ducky stimuli, both familiar and novel items were equally relevant to the observer, precluding the need for stronger attentional engagement by the familiar items. Secondly, if anything, familiarity would rather be associated with a reduction in attention, given that novel items are more salient and have stronger capacity to attract attention (Escera et al., 1998).

The experimental effects of familiarity and expectation had distinct time courses. The reduction in activity for familiar (compared to novel) items was apparent already 200 ms after the onset of the visual sequence, while the activity reduction for expected items only was visible after 500 ms. This is likely due to the fact that familiarity is already defined for the first image – which may have been seen before (familiar) or not (novel). With respect to expectation, the first image of a sequence could never be predicted; only after observing the first image, a prediction could be made about the subsequent images. This may explain why expectation effects are visible only later in the visual sequence. Apart from this, it is conceivable that activity modulation due to familiarity is the consequence of local changes in LOC (Kaliukhovich and Vogels, 2011; Vogels, 2016), while expectation suppression is implemented by feedback from regions higher up in the visual hierarchy (Friston, 2005) and potentially the hippocampus (Buckner, 2010; Hindy et al., 2016) which send predictions down to LOC and V1. This may explain why the modulation by familiarity may be observed earlier than the modulation by expectation.

Our data suggest that activity modulations by expectation may mainly be caused by surprise enhancement rather than expectation suppression, since unexpected images elicited significantly larger activity than unpredictable items, whereas expected images were not statistically different from unpredictable images in terms of neural activity. In line with this, Kaposvari et al. (2016) found that expectation suppression was of smaller magnitude than surprise enhancement, which opens the possibility that expectation suppression may be a weaker signal and therefore we did not observe it in our data. A recent study by Ramachandran et al. (2017), on the other hand, found strong evidence for expectation suppression and only limited evidence for surprise enhancement. As argued by these authors, the apparent discrepancy between these studies may relate to the rate of presentation of the visual stimuli. Rate of presentation in both our study and the study by Kaposvari et al. (2016) was markedly faster than in the study by Ramachandran et al. (2017), potentially resulting in a weaker phasic response to the image and thereby less potential for expectation suppression.

Interestingly, we observed an effect of expectation although many participants could not explicitly report the sequence for the predictable images. During debriefing, participants reported that they did not notice any specific order for the images. The behavioral assessment of sequence knowledge also showed that participants’ performance was on average near chance level when they were shown an image and had to report which image should follow. Nevertheless, the neural response showed a distinct difference between expected and unexpected conditions, suggesting that the neural effects of expectation are likely due to implicit predictions that occur outside the awareness of the observer. This is in line with previous studies showing that subjects can learn transitions without becoming aware of them (Reber, 1967; Clark and Squire, 1998; Alamia et al., 2016).

While we isolated neural effects of familiarity and expectation by independent manipulation, usually these concepts are heavily intertwined. In our everyday environments, we become familiar with images because they appear more often, which means that we also expect to see them more often. In this sense, familiarity can be viewed as one form of expectation. A noteworthy difference remains between the two, however: familiarity refers to the fact that the system has knowledge of certain past visual input, while expectation implies that the system is making predictions about upcoming visual information. In spite of this conceptual distinction, the neural consequences of these two processes appear similar in terms of neural activity modulation as well as the regions involved.

However, it is quite possible that the underlying neural mechanisms differ between familiarity and expectation. The time courses for the two manipulations differ, with a later modulation by expectation than by familiarity. Therefore, even though the resulting modulation in target regions appears rather similar, it is possible that the underlying sources are distinct. This could be the topic of future study using methods that allow to simultaneously map out the network of active regions, such as functional magnetic resonance imaging (fMRI).

## Acknowledgements

This work was supported by The Netherlands Organisation for Scientific Research (NWO Vidi grant 452-13-016 awarded to FPdL; NWO Vidi grant 864-14-011 awarded to JS; NWO Research Talent grant 406-13-001 awarded to PM, NWO Research Talent grant 406-16-525 awarded to MEM) and the EC Horizon 2020 Program (ERC starting grant 678286, ‘Contextvision’ awarded to FPdL).

